# Evolution and diversity of bat and rodent Paramyxoviruses from North America

**DOI:** 10.1101/2021.07.01.450817

**Authors:** Brendan B. Larsen, Sophie Gryseels, Hans W. Otto, Michael Worobey

## Abstract

Paramyxoviruses are a diverse group of negative-sense, single-stranded RNA viruses of which several species cause significant mortality and morbidity. In recent years the collection of paramyxoviruses sequences detected in wild mammals has substantially grown, however little is known about paramyxovirus diversity in North American mammals. To better understand natural paramyxovirus diversity, host range, and host specificity, we sought to comprehensively characterize paramyxoviruses across a range of diverse co-occurring wild small mammals in Southern Arizona. We used highly degenerate primers to screen fecal and urine samples and obtained a total of 55 paramyxovirus sequences from 12 rodent species and 6 bat species. We also performed illumina RNA-seq and de novo assembly on 14 of the positive samples to recover a total of 5 near full-length viral genomes. We show there are at least 2 clades of rodent-borne paramyxoviruses in Arizona, while bat-associated paramyxoviruses formed a putative single clade. Using structural homology modeling of the viral attachment protein, we infer that three of the five novel viruses likely bind sialic acid in a manner similar to other *Respiroviruses*, while the other two viruses from Heteromyid rodents likely bind a novel host receptor. We find no evidence for cross-species transmission, even among closely related sympatric host species. Taken together, these data suggest paramyxoviruses are a common viral infection in some bat and rodent species present in North America, and illuminate the evolution of these viruses.

**Importance:** There are a number of viral lineages that are potential zoonotic threats to humans. One of these, paramyxoviruses, have jumped into humans multiple times from wild and domestic animals. We conducted one of the largest viral surveys of wild mammals in the United States to better understand paramyxovirus diversity and evolution.

## Introduction

Emerging viruses, and zoonoses in particular, represent a severe threat to human health (1–3). Recently, the SARS-CoV-2 pandemic has emphasized the need for more research towards better understanding viral evolution in wildlife sources (4, 5). Most research on emerging viruses is concerned with disease in humans (6). However, for many zoonoses, humans represent a dead-end host (7). There are large knowledge gaps when it comes to viruses that circulate in wild mammal species, of which only a small fraction can infect humans and cause disease (8). Therefore, to understand the broader evolutionary and ecological context of potential zoonotic viruses before they jump into humans, additional research focused on a wide range of natural hosts is needed.

Paramyxoviruses (PMVs) are a large group of non-segmented, negative-sense RNA viruses in the order Mononegavirales, some of which cause devastating disease in humans and wildlife. Examples of important PMVs include measles and respirovirus (formerly human parainfluenza virus, a common cause of childhood respiratory disease). A number of PMVs have been involved in deadly zoonoses in humans in recent years, such as Hendra and Nipah (9–11). These outbreaks have attracted growing interest in zoonotic PMVs. Furthermore, increased sampling and improvements in sequencing technologies have revealed a large number of novel PMVs in wild mammals. Most sampling efforts have focused on rodents and bats due to their diversity, abundance, and the significance of their role as hosts for other viral zoonoses (12–17).

Despite the growing interest in these novel paramyxoviruses, there are still uncertainties regarding phylogenetic relationships of novel viruses with established genera due to small sequence fragments being used for many studies, as well as what criteria should be used to split taxa (18). The International Committee on Taxonomy of Viruses (ICTV) currently recognizes eight genera in the subfamily *Orthoparamyxovirinae* (Family: *Paramyxoviridae*). Three of these genera (*Henipavirus*, *Morbillivirus*, and *Respirovirus*) cause well-known diseases in humans and other animals, and have been studied in great detail. Two other genera, *Ferlavirus* and *Aquaparamyovirus* have only been found in reptiles and fish. *Salemvirus* currently only contains one virus which infects horses. This leaves two genera that contain an assortment of understudied but growing clades of mammal viruses. *Narmovirus* is a newly described paramyxovirus genus that contains three rodent-borne viruses and one tree shrew virus, with very little known about the biology of these viruses (19–22). Finally, the recently recognized genus *Jeilongvirus* currently contains seven viruses associated with the rodent families Cricetidae and Muridae, along with one bat virus (23, 24). The genus *Shaanvirus* has been proposed to contain recently discovered bat-associated paramyxoviruses (such as the one mentioned above) that contain two unique loci (Transmembrane (TM) and Short Hydrophobic (SH)) compared to bat-borne henipaviruses, however, currently this virus is classified as a *Jeilongvirus*.

Host receptor tropism is an important feature for understanding PMV evolution and zoonotic potential. PMVs across the subfamily *Orthoparamyxovirinae* contain a diverse set of receptor types for recognition and binding of host cells. The receptor locus is either described as H (for Hemagglutinin), HN (for Hemagglutinin / Neuraminidase) or G (for attachment glycoprotein). HN-bearing viruses use sialic acid to bind to host cells, while H and G-bearing viruses use surface cell proteins (25). Although the ORF that codes for these proteins always occurs in the same position of the genome (3’ of the L gene), the host receptors vary and sequence identity is low. However, structurally they share similarities and all contain six-bladed β-propeller domains (25). Due to limited sampling in these groups with appropriate whole genome sequences that contain the receptor ORF, little is known about the evolutionary direction and frequency of cell receptor tropism changes.

Regarding the evolution of PMVs in wild mammals, there are still many gaps in our knowledge. First, much of the focus of PMV detection has been done on old world species and on just a few widely distributed host species. Second, these viral surveys have largely relied on a small section of the viral polymerase gene, which can be amplified and sequenced with highly degenerate ‘universal’ primers. These fragments are useful for understanding basic phylogenetic relationships, but additional genome level data is needed to understand gene evolution across PMVs. Third, PMVs have a diverse array of host receptors they use to gain entry to cells, however the direction of these host receptor switches is still unclear. Finally, it is not clear how long these viruses have been associated with host lineages and how frequently cross-species transmission events are occurring.

We sought to comprehensively study PMVs in southern Arizona to add to our understanding of host range, prevalence, diversity, genome evolution, and host receptor usage. Since nothing is known about PMV diversity in North American small mammals, we decided to catch and sample as many species as possible. We used these sequence data to better understand the evolution of PMVs locally in our focal species, in addition to using previously sequenced reference PMV genomes to better understand the global evolution of PMVs.

## Methods

### Sample collection

From 2015-2018, we captured a total of 358 bats and rodents across 12 different locations with a perennial water source in southern Arizona. Bats from multiple species were captured in mist nets set over water sources, extracted from the nets and put in brown paper bags. Bats were kept in these bags for approximately 20 minutes during which time they were weighed. We next removed the bats, measured forearm length and collected oral and rectal swabs using a PurFlock 0.14” Ultrafine swab (Puritan, Guilford, Maine). Feces and urine were collected from the bag if present. All feces and urine samples were stored in a 0.5 mL buffer consisting of 1x PBS and 50% Glycerol. These samples were held on ice until returning to the lab where they were stored in a −80°C freezer.

Rodents were captured using Sherman traps baited with oats and seeds and left open overnight. After capture, rodents were transferred to a zip lock bag to take weight and obtain an approximate body and tail length measurement. Next, oral and rectal swabs were collected and stored into identical buffer as above, and hind foot length was measured. Feces and swabbed urine were further collected from the trap and bag if present.

The study was approved by the University of Arizona Institutional Animal Care and Use Committee permit #15-583. The permit from the Arizona Department of Game and Fish was numbered SP506475.

### Molecular characterization

In a biosafety level-2 laboratory, 2-ml centrifuge tubes containing buffer and swabs with either feces or urine were initially spun at 4,000xg for 5 min to pellet debris. The supernatant was transferred to a new 2-ml centrifuge tube and spun at 16,000xg for 1 min to pellet debris again. 200μL of this supernatant was used in the downstream extraction. Total RNA and DNA were extracted from fecal and urine samples using the QIAamp MinElute Virus Spin Kit (Qiagen, Valencia, CA, USA), but using 20μg of linear polyacrylamide (VWR, Radnor, PA, USA) per 200ul of sample instead of poly(A) RNA as a carrier to increase recovery of low quantities of nucleic acid and to not interfere with downstream high-throughput sequencing (26). DNA/RNA were eluted twice with 40μL of molecular grade water.

Reverse transcription was performed in 20μL final reaction volumes with the GoScript kit (Promega, Madison, WI, USA) and random hexamers (Thermo Fisher, Waltham, MA, USA). 8ul of extracted RNA/DNA were mixed with random hexamers (final concentration: 2.5μM) and dNTP (NEB, Ipswich, MA, USA). This volume was incubated in a heat block for 5 min @ 70°. Samples were chilled on ice for 1 minute and then a mixture containing 5x reaction buffer, 3mM MgCl2, 1μL Reverse Transcriptase, and 1μL of Recombinant Ribonuclease Inhibitor were added. Samples were put in a thermocycler at 25°C for 5 minutes and then at 42°C for 1 hour. Finally, reverse transcriptase was inactivated at 70° for 15 minutes.

cDNA was then screened using published primers targeting a conserved region of the polymerase gene of PMV from the Respiro-, Morbilli-, and Henipavirus (RMH) genera, or more specifically, viruses of the subfamily *Orthoparamyxovirinae* (27). Hemi-nested PCR was performed with Taq DNA Polymerase (NEB, Ipswich, MA, USA). First round PCRs were performed with 5μL of cDNA, and a final concentration of 1uM of each primer, and 2.5mM MgCl. PCR steps included 1 min @95°C, 40 cycles of 95°C for 30 seconds, 50°C for 30 seconds, and 68°C for 1 minute, followed by a final extension at 68°C for 5 minutes. Second round PCRs were identical with the exception of using 1μL from the first round PCR as input. To obtain more sequencing information following successful amplification and sequencing of RMH amplicons, an additional degenerate primer set (PAR) spanning ~530 nt was used on the positive samples, amplified, and sequenced (27). PCR conditions were the same as above. All reactions were visualized in a 1.5% Agarose gel containing 1X GelRed fluorescent nucleic acid dye (Biotium, Fremont, CA, USA). Positive samples were sequenced on an Applied Biosystems 3730XL DNA Analyzer at the University of Arizona Genetics Core.

For whole PMV genome sequencing via Illumina technology, library prep was performed on extracted DNA/RNA with the Trio RNA-seq kit (NuGEN, San Carlos, CA, USA). Sequencing was performed at the DNASU sequencing core of Arizona State University on the NextSeq 500 using the paired end mid-output kits with either 2×75 or 2×150 read lengths.

Two of the samples that were subjected to Illumina sequencing (BA5 and BC5) were incomplete, and an iterative process of targeted PCR amplification followed by Sanger sequencing was performed to fill in gaps between contigs produced by illumina sequencing. We modified the reverse-transcription protocol to deal with secondary structure, which caused our initial attempts to fail. Instead of random hexamers we used the forward primers at a final concentration of 0.5μM for each sequence to initiate reverse transcription (Primers used are given in Supplementary Table 1). We also included 1μL of DMSO (VWR, Radnor, PA, USA). Due to higher incubation temperatures we used 1 μL RNasin Plus Ribonuclease Inhibitor (Promega, Madison, WI, USA). All other methods were identical to previous reverse transcriptions with the exception of changing the main incubation step to 55°C for 1 hour in order to disrupt RNA secondary structure. Following the RT step, cDNA was incubated with 10U Ribonuclease H (ThermoFisher, Waltham, MA, USA) for 20 minutes at 37°C. First round PCRs were done with 5μL of cDNA, 0.4uM final concentration of each primer, and the Takara LA HS Taq long range polymerase (Takara, Mountain View, CA, USA). PCR conditions used were 94°C for 3 minutes, followed by 40 cycles of 98°C for 10 seconds, and different extension times and temperatures depending on the specific amplicon, with a general rule of 1 minute per kb of amplicon.

Finally, on all complete genomes we attempted 3’ and 5’ RACE using custom methods for PMVs based on published work (28). All attempts were unsuccessful and so the 3’ and 5’ ends remain unmapped.

### Host molecular identification

All mammals were initially identified to species in the field using metrics such as forearm length and weight in bats, and hind foot length in rodents. For any specimen for which morphological species identification was unsure, and for a general subset of animals, we used a well-established published *cytochrome b* (*cytB*) gene barcode PCR to determine species ID (29). In no case did the *cytB* barcode sequence disagree with our initial identification. However, for the established species *Myotis californicus* and *Myotis ciliolabrum* the taxonomic status is unclear; with neither the *cytB* locus or external morphological traits able to discriminate the species (30). Therefore, in all following descriptions we refer to these samples as coming from “Myotis ciliocal” by lumping them together.

### Sequence analysis

Sanger sequences were assembled from forward and reverse chromatograms in Geneious V8.1.5 (Biomatters, Auckland, New Zealand). Primer sequences were trimmed and chromatograms were visually inspected to ensure accurate base calling. For cases where chromatograms had low quality peaks, we resequenced the amplicon to attempt to get higher sequence quality.

Raw reads were de novo assembled with Trinity v2.6.5 using default settings and trimming enabled (31). All assembled contigs were then compared with proteins from *Jun jeilongvirus* (J virus) (Accession:YP338075.1-YP_338085.1) with tblastn and default parameters (32). Contigs that matched J-virus with e-values less than 1×10^−5^ were then compared to the complete non-redundant GenBank nucleotide database using BLAST to ensure each hit did not have a more significant match to something other than a PMV. All RMH, PAR, and cytB sequences have been deposited in GenBank (Accessions XXX-XXX).

### Phylogeny construction

Alignments for each ORF were first aligned in Geneious v8.1.5 (Biomatters, Auckland, New Zealand) using the Translation Align feature, and then trimmed with an online server running Gblocks v0.91b to remove uninformative/highly variable regions from the alignments (33). The third codon site was stripped from all alignments due to high sequence saturation. Phylogenies were made in IQTREE v2.0 (34) with automatic model selection and 1000 bootstrap replicates.

### Homology modeling

The H/HN/G locus from sample BC5 was used as a query sequence for structural homology modeling on the online Swiss-model server (35). The structural alignment with the best QMEAN score of −3.41 and a GMQE score of 0.53 was the template 1V3D with 32.9 % sequence identity. 1V3D is the previously determined crystal structure of the Haemagglutinin-neuraminidase protein from Human Parainfluenza Virus Type III, a paramyxovirus of the genus Respirovirus (36). To visualize the active site comparison between the modeled BC5 structure with 1V3D we used Chimera v1.13.1 (37).

### Codivergence

To assess whether PMVs and bats share a history of codivergence, we downloaded *cytB* references from GenBank and used them to construct a host phylogeny. We then tested whether the host/virus phylogeny shared a similar typology than would be expected by chance using the program parafit implemented in R (38).

### Statistical analysis

To test whether positivity rate varied by mammalian family we tested the effect of family on positivity using a binomial general linear model implemented in R with the package lme4 (39).

## Results

### Paramyxovirus detection and sequencing

Fecal and urine samples from a total of 358 individuals from 15 rodent and 18 bat species captured across southern Arizona were included in this study (Fig. 1). None of the animals we sampled showed overt signs of disease.

**Figure 1.**
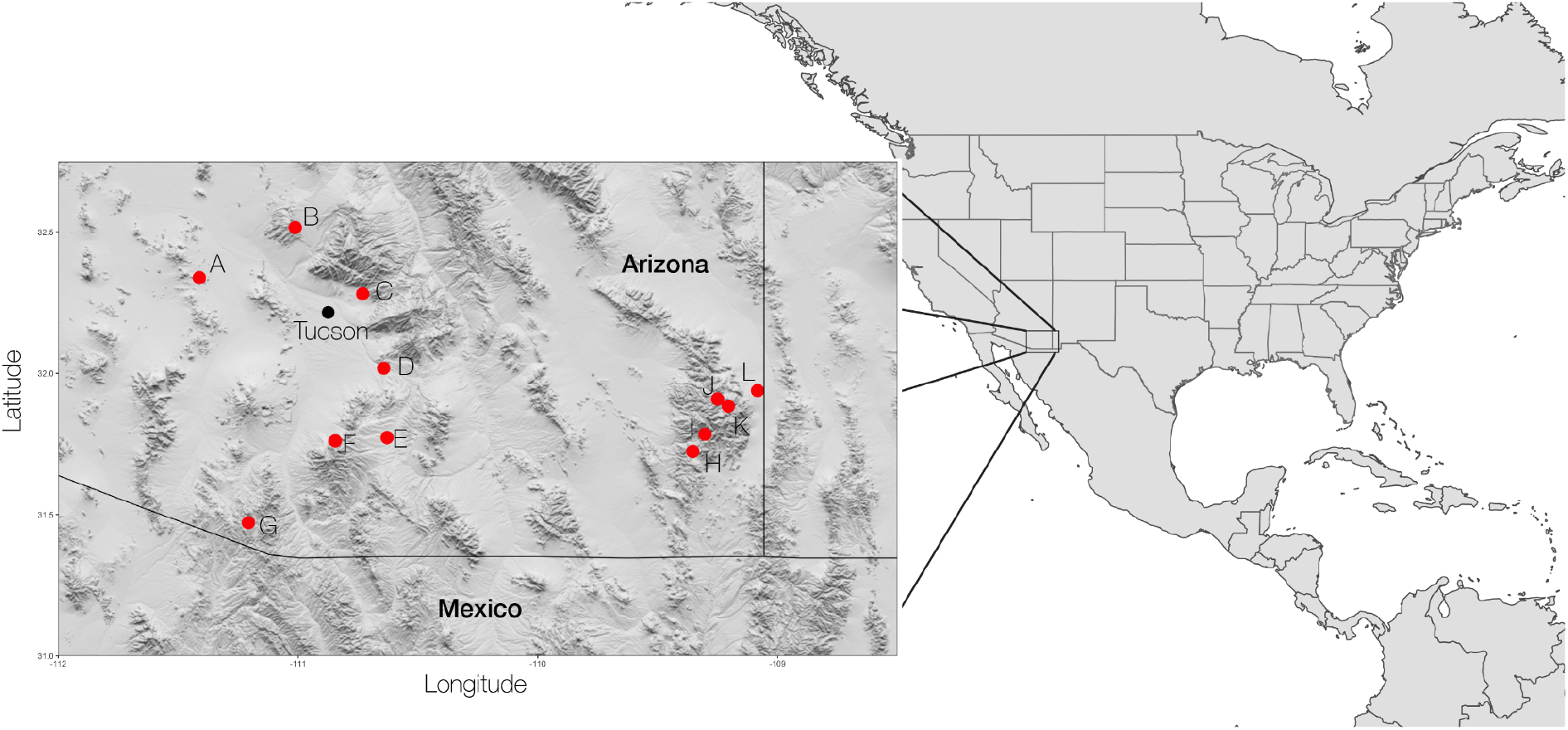
Elevation relief map showing sampling locations across SE Arizona. Each sampling location is represented with a red dot and a 1-letter code which corresponds to the information in Supplementary Table 1 with capture information for all species.

Samples were screened with hemi-nested degenerate primers previously designed to amplify viruses from the *Respirovirus, Henipavirus*, and *Morbillivirus* (RMH) genera within the subfamily *Orthoparamyxovirinae*. Overall, 53 individual samples were positive by the RMH primer set and a ~490 nt fragment was Sanger sequenced. 36% of all rodents tested were PMV positive compared to 9% of bats. The overall positivity rate of PMVs detected varied significantly between/among rodent and bat families even when accounting for sampling biases (p<0.001) (Table 1). Of the 53 RMH positives, 43 were also positive for a longer sequence fragment (PAR) and were Sanger sequenced. Two samples were positive only for the PAR primer set, bringing the total number of positive samples to 55. Information about all positive animals and their corresponding sequence ID are summarized in Supplementary Table 2.

**Table 1.** Detection of PMV by Mammal Order, Family, and Species.

| Rodentia |  |  |  |  |
| --- | --- | --- | --- | --- |
|  | Heteromyidae | Virus Positive | Tested | Percent Positive |
|  | *Dipodomys ordii* | 3 | 8 | 38% |
|  | *Dipodomys merriami* | 2 | 17 | 12% |
|  | *Dipodomys spectabilis* | 0 | 1 | 0% |
|  | *Chaetodipus baileyi* | 3 | 19 | 16% |
|  | *Chaetodipus penicillatus* | 2 | 5 | 40% |
|  | Cricetidae |  |  |  |
|  | *Peromyscus eremicus* | 1 | 1 | 100% |
|  | *Peromyscus leucopus* | 2 | 2 | 100% |
|  | *Peromyscus maniculatus* | 6 | 13 | 46% |
|  | *Peromyscus boylii* | 3 | 5 | 60% |
|  | *Neotoma albigula* | 2 | 6 | 33% |
|  | *Sigmodon arizonae* | 3 | 4 | 75% |
|  | *Reithrodontomys megalotis* | 4 | 4 | 100% |
|  | *Reithrodontomys fulvescens* | 0 | 1 | 0% |
|  | *Reithrodontomys montanus* | 0 | 1 | 0% |
|  | *Onychomys torridus* | 1 | 3 | 33% |
|  | Rodent Total | 32 | 90 | 36% |
| Chiroptera |  |  |  |  |
|  | Vespertilionidae |  |  |  |
|  | *Lasiurus blossevillii* | 0 | 1 | 0% |
|  | *Lasiurus cinereus* | 0 | 4 | 0% |
|  | *Lasiurus xanthinus* | 0 | 16 | 0% |
|  | *Myotis ciliocal* | 5 | 12 | 42% |
|  | *Myotis auriculus* | 0 | 7 | 0% |
|  | *Myotis thysanodes* | 0 | 5 | 0% |
|  | *Myotis yumanensis* | 0 | 22 | 0% |
|  | *Myotis velifer* | 2 | 67 | 3% |
|  | *Myotis volans* | 0 | 3 | 0% |
|  | *Antrozous pallidus* | 4 | 21 | 19% |
|  | *Eptesicus fuscus* | 0 | 22 | 0% |
|  | *Corynorhinus townsendii* | 0 | 2 | 0% |
|  | *Parastrellus hesperus* | 0 | 11 | 0% |
|  | Phyllostomidae |  |  |  |
|  | *Leptonycteris yerbabuenae* | 0 | 10 | 0% |
|  | *Choeronycteris mexicana* | 0 | 2 | 0% |
|  | *Macrotus californicus* | 1 | 7 | 14% |
|  | Molossidae |  |  |  |
|  | *Tadarida brasiliensis* | 7 | 45 | 16% |
|  | *Nyctinomops femorosaccus* | 4 | 11 | 36% |
|  | Bat Total | 23 | 268 | 9% |
|  | Overall | 55 | 358 | 15% |

For 3 of the 14 deep-sequenced samples (AK5, M14, AB1) we recovered near full-length PMV genomes after de novo assembly of the reads. Two additional samples had >50% coverage and were filled in with subsequent Sanger sequencing (BC5, BA5). Finally, one sample had <10% coverage but we were able to recover near full-length sequences for the L gene (BK1). The remaining samples had reads that covered <5% of the PMV genome and were not used for the remainder of analyses.

### Phylogenetic relationships among novel and known paramyxoviruses

The maximum likelihood phylogeny of the 43 concatenated RMH/PAR sequences characterized in this study together with representative sequences from all *Orthoparamyxovirinae* genera is shown in Figure 2. The novel RMH/PAR sequences fell into 3 clades, which are described below.

**Figure 2.**
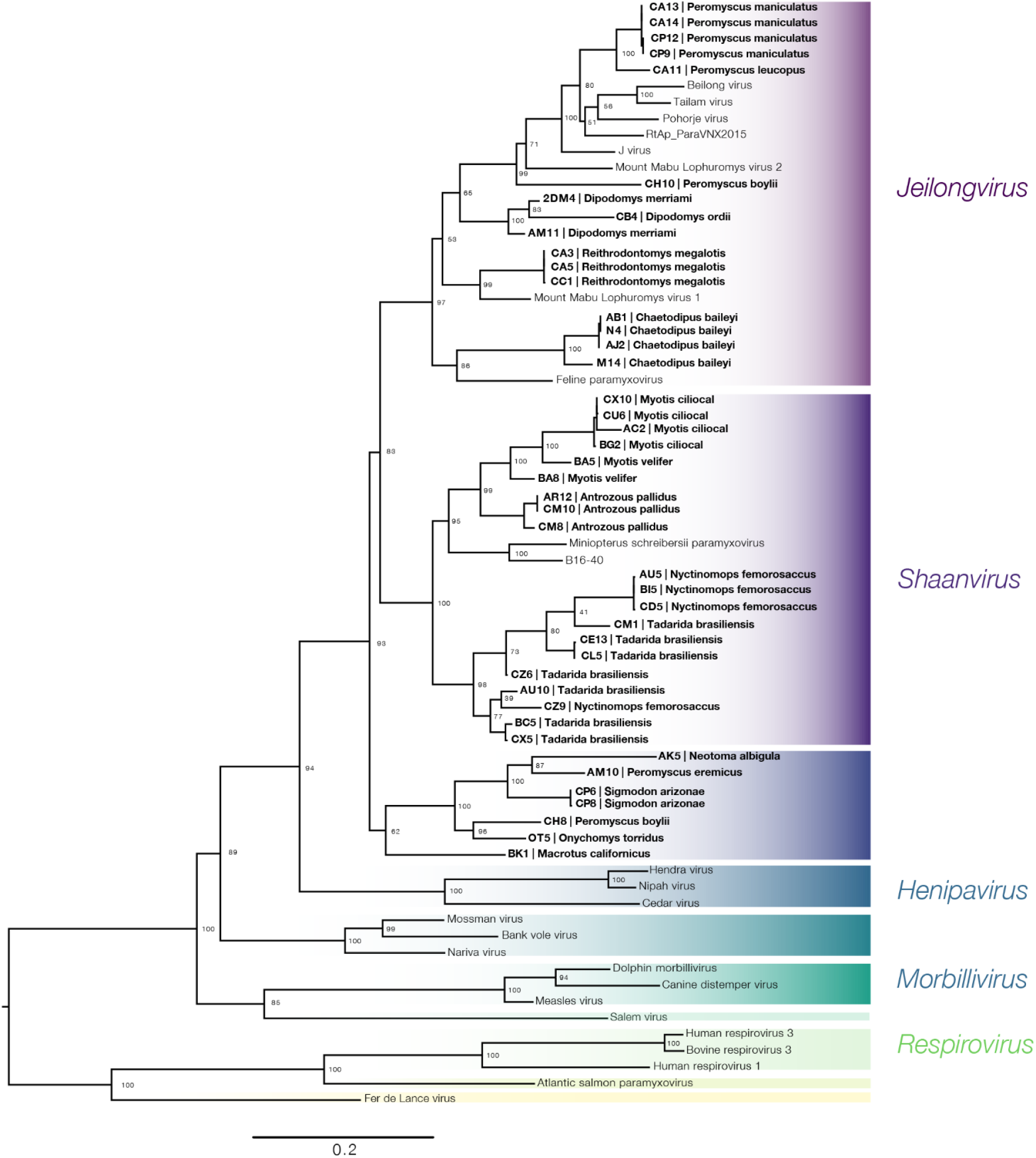
Maximum likelihood phylogeny from the concatenated RMH (~490 nt) and PAR (~550 nt)_sequences with reference paramyxoviruses and novel paramyxoviruses found in this study (bold). Every recognized or proposed genus within *Orthoparamyxovirinae* are colored, with major clades named. Tree was midpoint rooted.

Sequences from 19 out of 20 RMH/PAR positive bat samples from this study formed a clade with other bat viruses in the proposed genus *Shaanvirus*, which has not yet been accepted by the ICTV, but is a tentative genus of PMVs from bats that contains transmembrane (TM) and short hydrophobic (SH) loci, which distinguishes them from other bat PMVs in the genus *Henipavirus* (24).

Additionally, 15 RMH/PAR positive samples collected from the rodent families Cricetidae and Heteromyidae fall in a well-supported clade (bootstrap = 97), which contains other viruses from the recently recognized genus *Jeilongvirus*. Additionally, seven of our samples (one from the bat species *Macrotus californicus* and six from Cricetidae rodents) form a poorly supported clade (bootstrap = 62) distinct from any previously reported PMV genus.

However, because we recovered longer stretches of sequence for two of the samples in this clade (a near full-length genome (AK5) and a near complete coding sequence of L (BK1)), we were able to test whether the addition of further sequence information would change the placement of these samples. A maximum likelihood phylogeny based on the alignment of the two most conserved genes, M and L, placed AK5, a rodent PMV, with the other *Jeilongvirus* sequences (but with low support; bootstrap=70) (Supplementary Figure 1), and not the original placement as a novel clade. Furthermore, a phylogeny of the near full-length L gene placed the bat genome BK1 with other *Shaanviruses* (bootstrap=71) (Supplementary Figure 2). This contradicts the previous phylogeny based on the shorter RMH+PAR sequences and suggests the above mentioned novel clade of rodent and bat PMVs is likely an artifact of low information content of short sequences, and that all RMH/PAR sequences from this study are associated with established or proposed genera (*Jeilongvirus, Shaanvirus)*.

Phylogenetic analysis of the larger dataset from an alignment of all 53 RMH sequences (490 nt) showed that two samples from the rodent family Cricetidae contained viruses (BA15, from *Neotoma albigula* and CC3, from *Reithrodontomys megalotis*) clustering with reference viruses from the *Narmovirus* clade (Supplementary Figure 3). However, this clade may again be an artefact due to the short sequences used in this phylogeny.

Gene trees from the loci N, M, H/HN/G, and L show different relationships between genera depending on the gene analyzed (Figure 4). Overall branch support and groupings across loci are low. Notably, the phylogeny of the attachment protein locus H/HN/G shows a sister relationship between *Respirovirus* and all other *Jeilongvirus* and *Shaanvirus* sequences for the attachment protein locus, despite *Respirovirus* being the outgroup lineage for all other loci.

**Figure 3.**
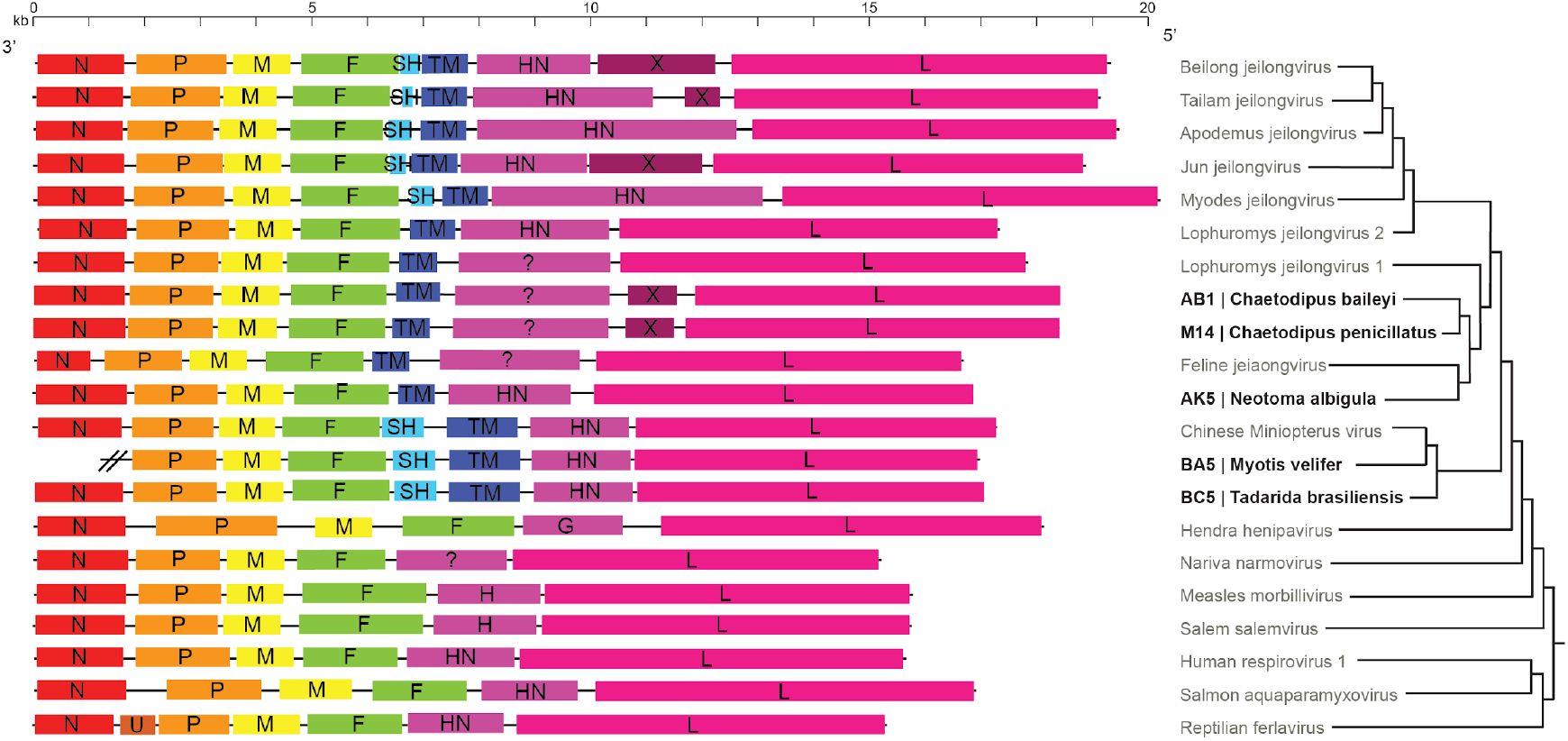
Genome structure with putative genes based on ORF homology of reference paramyxoviruses (gray) with novel viruses from this study (in bold). The phylogeny on the right is based on amino acid alignments of complete L genes that were cleaned using G-blocks to only include amino acid positions where homology could be reliably assessed. The phylogeny is only meant to show the putative relationships of the viruses to understand the gain and loss of different genes. Branch lengths are not to scale and no node support was included. Question marks are used on the H/HN/G locus if the gene does not contain the canonical sialic acid binding receptor and has a putative unknown receptor. The BA5 assembly did not include the most 3’ end of the genome and so is missing.

**Figure 4.**
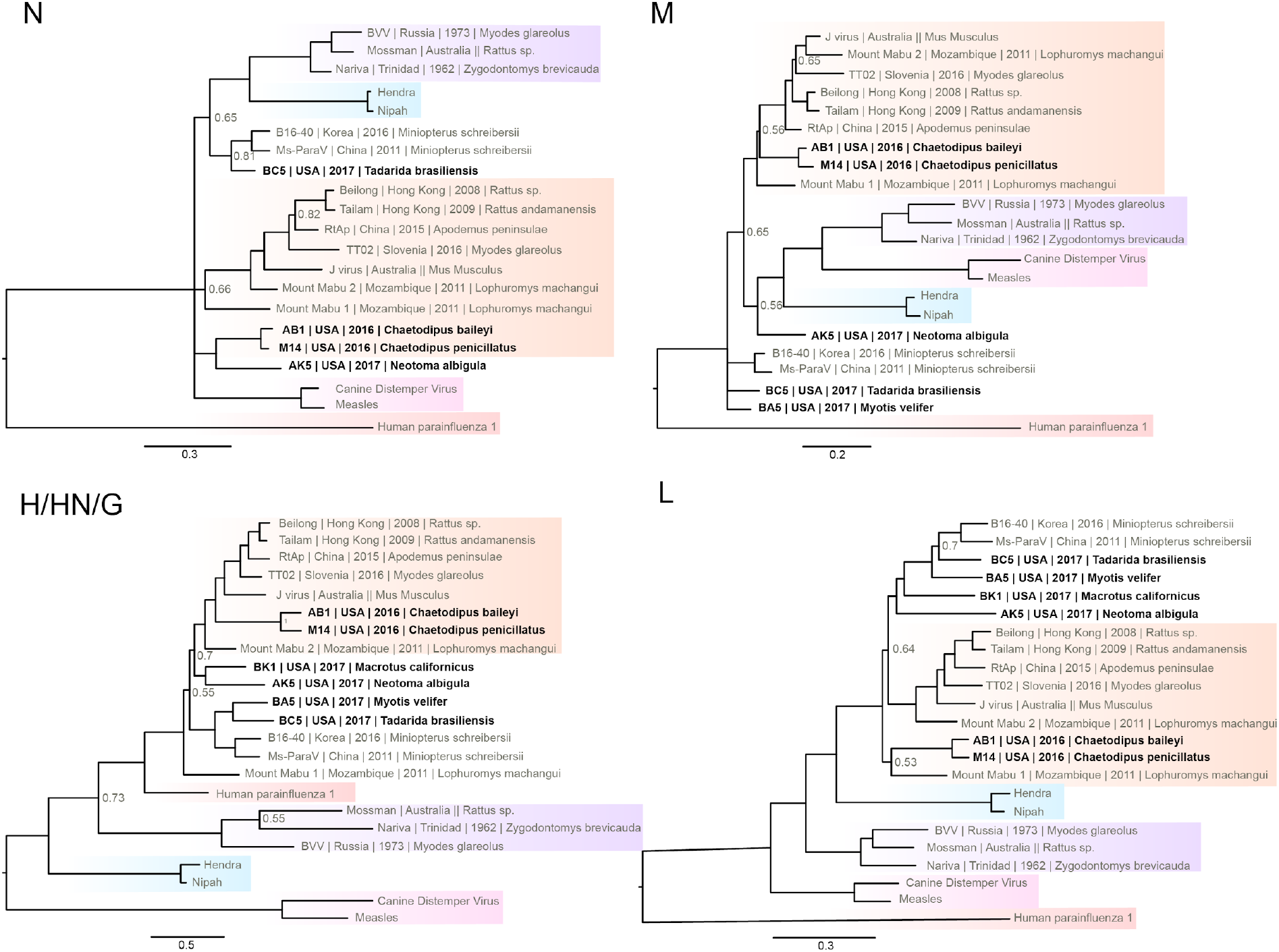
Maximum likelihood gene trees built from complete sequences for four genes from representative paramyxoviruses for which complete genome sequences are available (in bold, from this study). Established genera are colored red, light blue, pink, and purple for R*espirovirus, Henipavirus, Morbillivirus, and Narmovirus*, respectively. Only bootstrap values <0.85 are shown. For each gene tree, sequences were clipped with Gblocks, and 3rd position in codon stripped. Trees were midpoint rooted.

### Genome organization

We attempted to recover the terminal 3’ and 5’ ends of the virus genomes using previously published RACE techniques, but were unable to amplify any sequence, perhaps due to low amounts of template (28). However, apart from missing the N gene in BA5, we sequenced the complete coding regions for five near full-length genomes from our original sample size of 14.

The open reading frames of the five near full-length genomes along with representative genomes from other *Orthoparamyxovirinae* genera are shown in Figure 3. The coding genome lengths, starting and ending at the predicted open reading frames, from the *Jeilongviruses* AB1 (collected from *Chaetodipus baileyi)* and M14 (from *Chaetodipus penicillatus)* are 18,495 and 18,471 nt long, respectively. The coding genome of AK5 (collected from *Neotoma albigula)* is shorter, at only 16,182 nt. BC5 and BA5, from the bat species *Tadarida brasiliensis* and *Myotis velifer*, respectively, shared an identical genome structure to previously described full-length bat PMVs from Asia in the putative genus *Shaanvirus*. The BA5 assembly is missing the 3’ end of the genome which includes the N gene due to low coverage, therefore the total length of the sequenced genome is 15,319 nt from just before the start of the P gene to the end of the L gene. BC5, a *Shaanvirus* from the host species *Tadarida brasiliensis*, is a total of 17,150 nt.

The *Jeilongviruses* AB1, M14, and AK5 all have a predicted ORF that contains a transmembrane domain (TM) but lack an ORF that corresponds to the short hydrophobic (SH) gene, which is found in some, but not all *Jeilongviruses*. The two novel *Shaanvirus* bat PMVs have two ORFs that likely correspond to SH and TM genes also found in previously reported *Shaanviruses*. BA5 and BC5 SH ORFs are 233 and 245 amino acids long, similar to 239 amino acids for the previously reported Asian bat virus B16-40. These SH ORFs are longer than the 69 amino acids for *Jun jeilongvirus* (J virus).

The TM protein of BA5 and BC5 are 442 and 418 amino acids long respectively. The TM of the Asian bat virus B16-40 is 561 amino acids. In all three cases the bat TM protein is much longer than the TM protein of rodent PMVs in *Jeilongvirus*. For example, AB1 and M14 TMs are both only 259 amino acids long. Sequence similarity of the TM locus is very low across sequences. Comparing the bat PMVs BC5, BA5, and the previously published Asian bat PMV B16-40, the average pairwise amino acid identity was 16.5%. The average pairwise amino acid identity at the TM locus for the three bat PMVs compared to the *Jun jeilongvirus* virus was 5%.

### Comparisons of the host receptor binding site of HN from novel Paramyxoviruses to Respiroviruses

We further explored the relationship and function of the H/HN/G protein among our novel viruses and known representative sequences from the other known *Orthoparamyxovirinae* genera. Structural homology modeling showed a high structural similarity between the bat PMV BC5 H/HN/G protein and the experimentally determined crystal structures of the HN protein from Human Parainfluenza 3 (Now known as Human Respirovirus) (PDB:1V3D) (Figure 5) when bound to human sialic acid (a sugar used as a receptor for cell binding). All hydrogen bonds observed between the HPI3 binding site and the sialic acid ligand N-Acetylneuraminic acid (Neu5Ac) were predicted to occur in BC5, strongly suggesting BC5 likely uses a similar sugar to bind host cells. Furthermore, the residues involved in the active site binding in HPI3 were shown to be largely conserved in viruses from the putative *Shaanvirus* and *Jeilongvirus* genera with the exception of our samples AB1 and M14, and the reference genomes *Lophuromys jeilongvirus 1* and *Feline jeilongvirus* (Figure 5). AB1 and M14 were both collected from Heteromyid rodents, while the three other genomes we sequenced were from Cricetid rodents and bats. Therefore, the two Heteromyid PMVs likely have H/HN/G proteins with a novel divergent undiscovered co-receptor. These results show that despite low sequence identity, three of the five novel whole genome sequences we recovered likely bind a sialic acid in a similar fashion as *Respiroviruses* and other previously described rodent and bat PMVs.

**Figure 5.**
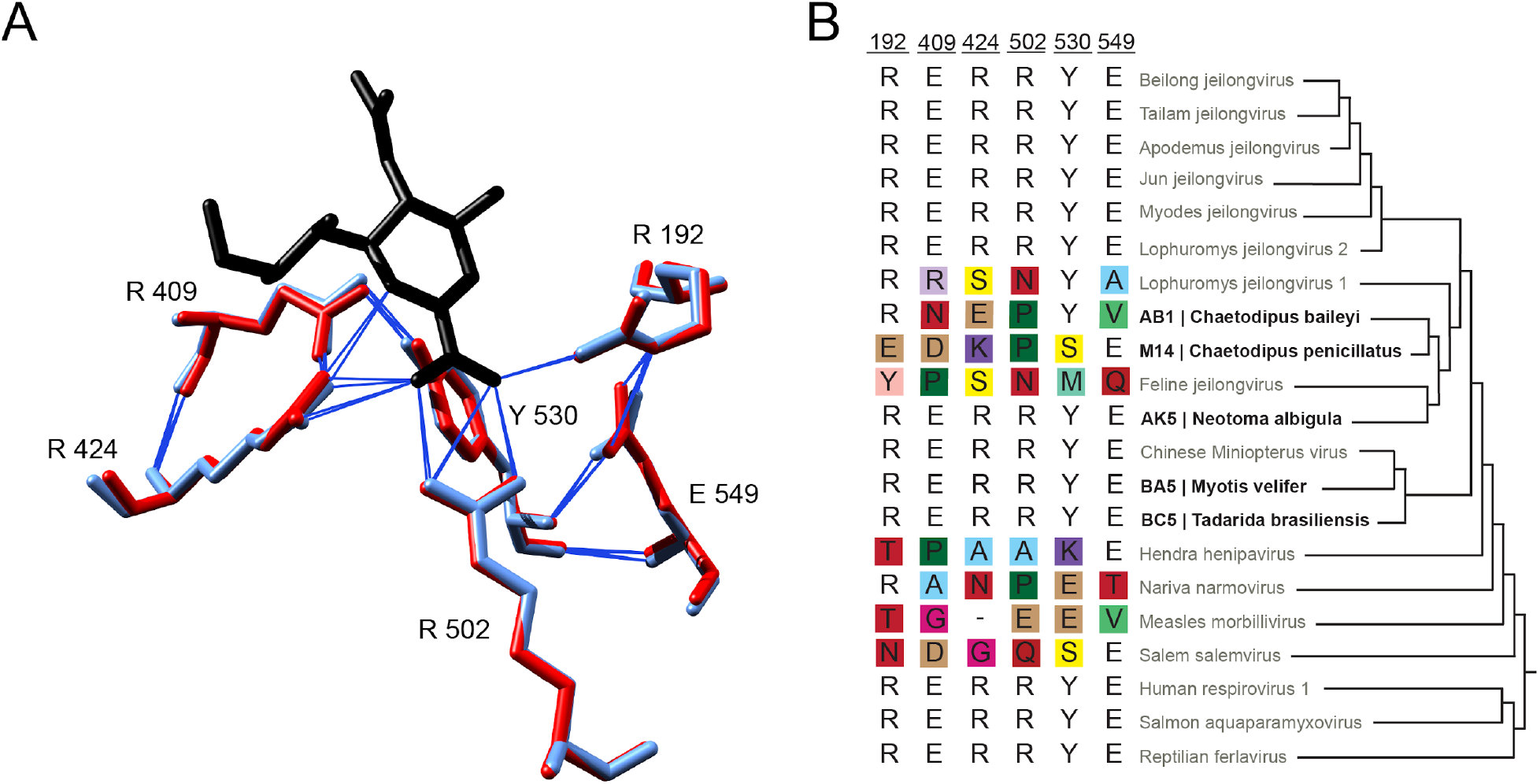
Comparison of key amino acids in the active site of sialic acid binding viruses at the HN locus. (A) shows the positions of the binding site amino acid residues and numbering of Human parainfluenza-3 (now Human respirovirus), PDB:1V3D, in red. Residues in light blue are the inferred binding site of BC5 (from *Tadarida brasiliensis)* based on structural modeling. In black is the sialic acid ligand N-Acetylneuraminic acid, and blue lines represent inferred hydrogen bonds between residues and the sialic acid. (B) shows the same active site amino acid residues involved in binding for different representative paramyxoviruses (light gray) with the novel paramyxoviruses from this study (in bold). Amino acid differences from the sialic acid binding sites are colored. The phylogeny on the right is the same as in Fig. 2.

### Host specificity

To test the amount of host specificity in PMVs, we choose to sample an area where two closely related mammalian sister species occur in relatively high densities. *Peromyscus maniculatus* and *Peromyscus leucopus* are closely related species in the rodent family Cricetidae that likely diverged ~500,000 years ago and still experience introgression in parts of their range (40). Over two nights of trapping in two separate areas where both species occur together, we captured eleven *P. maniculatus* and two *P. leucopus*. From these, eight of the *P. maniculatus* were positive by PCR for paramyxovirus and both of the *P. leucopus* were positive. Monophyly of these viruses were observed with respect to the host species (Figure 6).

**Figure 6.**
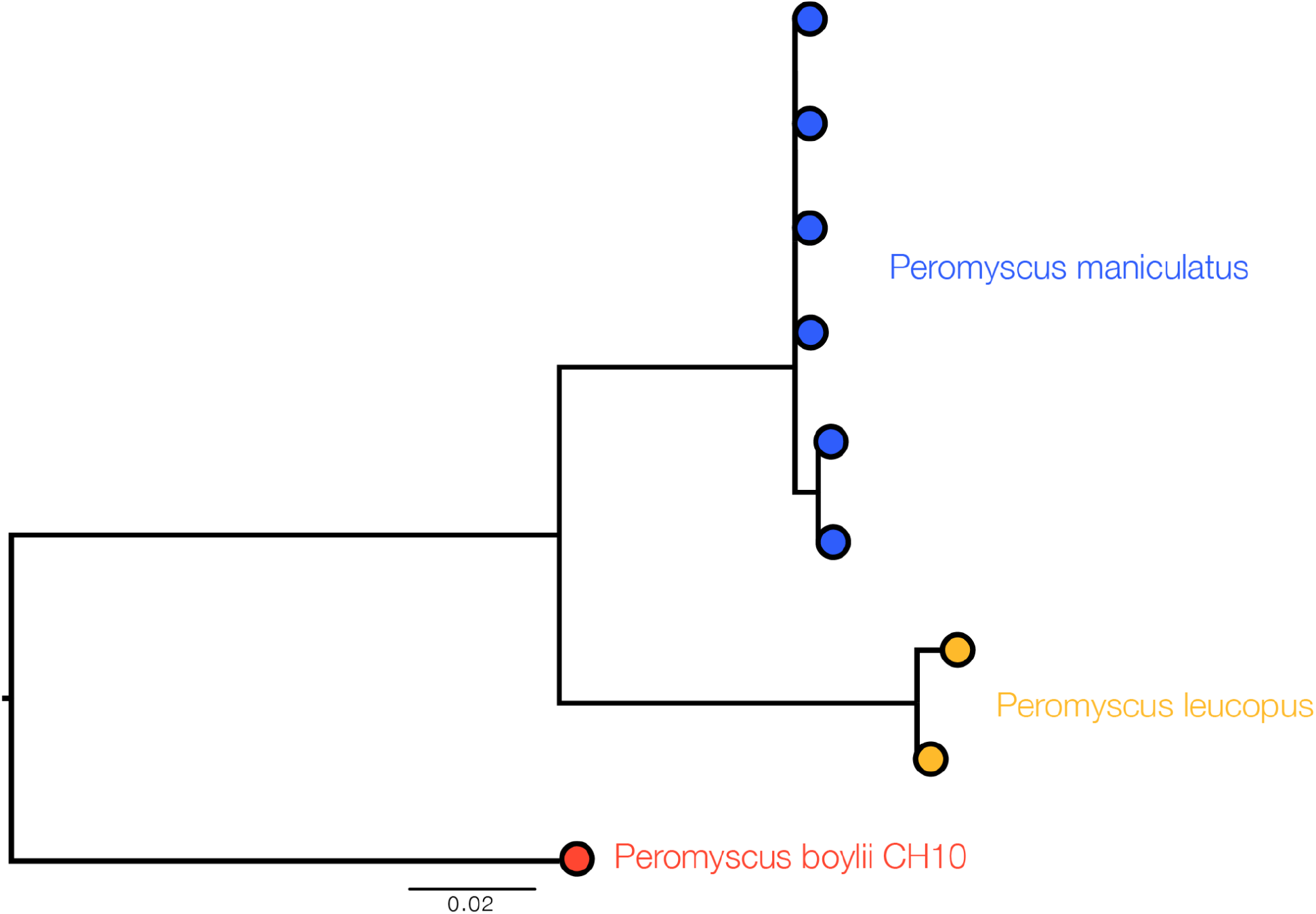
Maximum likelihood RMH phylogeny showing relationship of *P. maniculatus* and *P. leucopus* paramyxoviruses sampled in a single high density area. Tree is rooted with the closest Peromyscus paramyxovirus we sampled (sample CH10, from *P. boylii*)

Due to multiple circulating PMV clades in rodents, we focused our co-divergence analysis on bat PMVs from the putative genus *Shaanvirus* and compared them to host trees (Figure 7). The association between PMV and host is highly significant (p<0.001). Notably, some bat PMV sequences do not form clades relative to host species (for example, viruses from *Tadarida brasiliensis)* with respect to host species, suggesting processes such as host switching and within-host duplication events have also occurred during PMV evolution.

**Figure 7.**
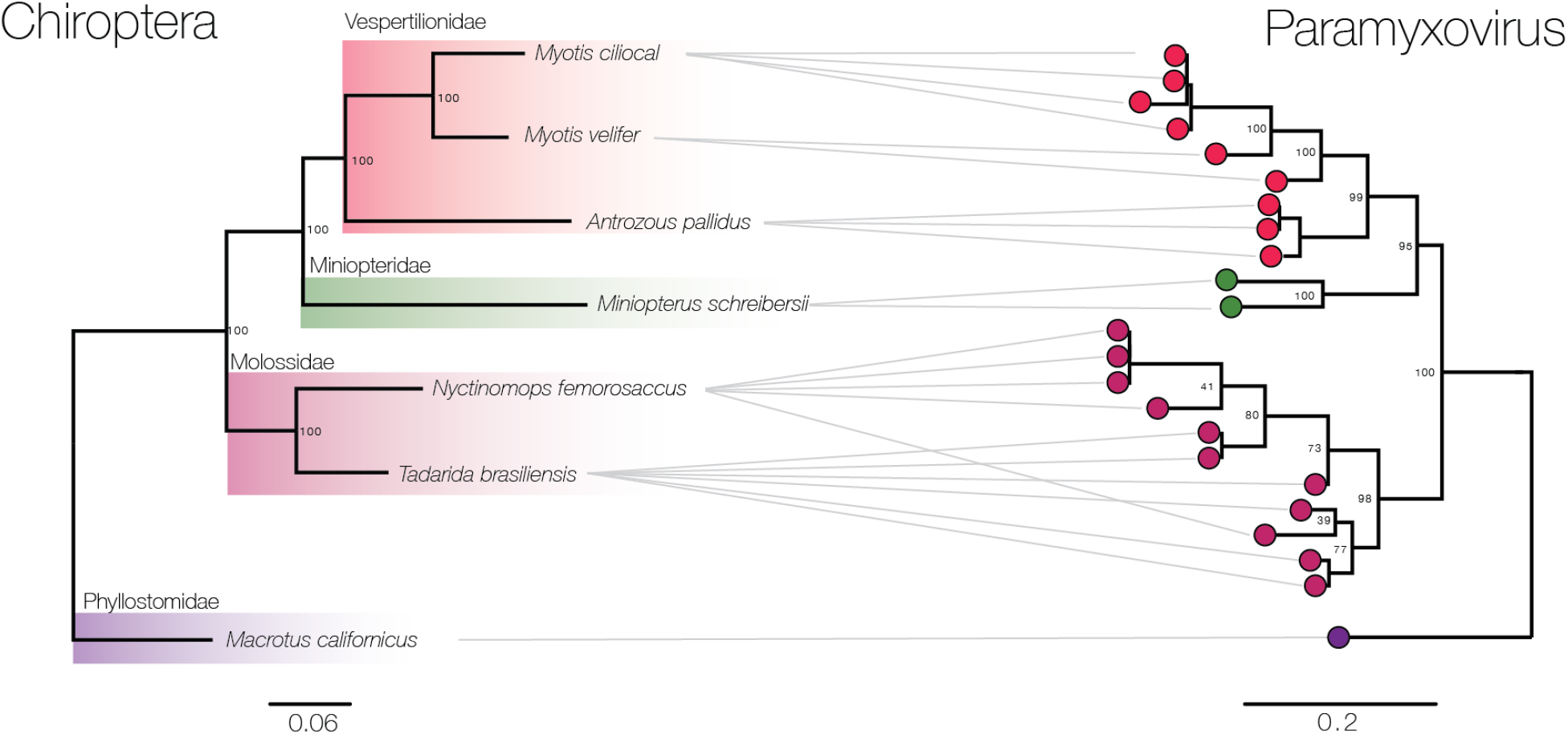
Maximum likelihood phylogenies showing relationships between host (left) and virus (right) of putative *Shaanvirus* members. Bat families are shaded.

## Discussion

Here, we add to the growing knowledge of the biology of bat and rodent associated PMVs. We characterized these new North American PMVs and placed them in context with previously described PMVs using a combination of phylogenetic analyses, whole genome sequencing, and structural homology modeling of the viral/host binding interface.

We detected an overall prevalence in 36% of rodents and 9% of bats, which is largely consistent with other studies which detected PMV prevalence of 25% in rodents of Madagascar (17), 3% in bats sampled in 15 locations across the world (12), 21% of rodents in Zambia (41), 1% of bats in Italy (42), and 4% of bats on islands of the southwest Indian Ocean (17). In general, it appears that rodents support a higher overall PMV infection burden than bats. The high infection burden in rodents could be due to chronic infections of PMVs, something that has been observed for other rodent-borne viruses, such as hantaviruses (43). Previous work has shown there is a seasonality pattern to PMV shedding in bats, however we did not have enough data within and across years to do a similar analysis (12). The observed lower prevalence in bats could thus possibly also be explained by sampling in low-prevalence seasons, which in turn would imply an absence of such PMV seasonality in rodents.

In our study we did not attempt to discern if these novel PMV are pathogenic to their natural hosts, or if they could possibly cause disease in humans. None of the animals we sampled showed overt signs of disease. Studies on similar novel PMVs from bats and rodents in other studies do offer some insight on the pathology of paramyxoviruses in wild mammals. When *Jun Jeilongvirus* was first isolated in 1977 in Australia it came from moribund *Mus musculus* (23), however later studies did not observe any disease in infected mice (44). An additional J virus from *Mus musculus* was isolated with only a few nucleotide differences from the original sequence which did cause severe disease in mice (45). Furthermore, pathology was noted in the kidney of an infected pipistrelle bat (13). Overall, pathology may be limited to specific host/virus interactions and more work needs to be performed regarding determinants of PMV virulence in wild mammals.

The difficulty of performing rigorous phylogenetics on extremely divergent viruses such as PMVs from different genera is highlighted by the phylogenetic discordance found depending on the gene used. Phylogenetic discordance also emerges even when using different fragments within the same gene. The ~1kb RMH/PAR fragment which is commonly used in studies as it is reliably amplified by degenerate primers is within the L gene. In this fragment, AK5 was found to be sister to all other bat and rodent PMVs from this study (Fig. 3). Although recombination could produce this pattern, recombination is thought to be rare in negative sense RNA viruses (46). However, when we used a much longer sequence (4,869 nt), of the concatenated M and L loci, AK5 was sister to all other *Jeilongviruses* with moderate branch support. Phylogenetic relationships should thus be interpreted with caution when using short fragments of conserved diverse viruses, and should not be used for taxonomic classification.

All PMVs share the same 6 core genes (N, P, M, F, (H/HN/G), and L). In recent years new genes have been discovered that add to the complexity of PMV genome content. The short hydrophobic (SH), transmembrane (TM), and unknown (X) have all been detected in various bat and rodent PMVs. Although knowledge on the function of these genes is gradually emerging, the evolutionary history remains unclear (47). From a phylogeny based on the complete amino acid sequence of L, which is the longest and most conserved of all PMV genes (Fig 2; right side), it appears that SH and X have multiple origins. Although bat SH proteins contain a hydrophobic domain, and bat TM proteins contain a transmembrane motif, and are in the same respective genome positions as *Jeilongvirus* SH and TM loci (between F and H/HN/G), it is not clear whether SH and TM from *Shaanvirus* are homologous with rodents SH and TM. Bat PMV SH and TM are significantly longer and in all cases share extremely low sequence similarity with *Jeilongvirus* SH and TM, approaching what would be the expected identity if two random amino acid sequences were aligned. More research is needed to understand the function of TM and SH in novel bat and rodent PMVs. What is clear however, is that bat PMVs such as BA5, BC5, and the Asian Miniopterus sequence B16-40 are not only phylogenetically distinct from rodent *Jeilongvirus* but may also each contain non-homologous proteins, and should likely be classified in their own genus of bat-borne PMVs that contain unique SH and TM ORFs compared to rodents, in the previously proposed genus *Shaanvirus (24)*.

PMVs exhibit remarkable receptor tropism flexibility throughout the course of their evolution. Across the entire breadth of PMV entry strategies, the most common appears to be a respirovirus-like attachment to N-Acetylneuraminic acid. However, within the overall phylogeny there are clear exceptions to this case, and these should be studied further. For example, *Morbillivirus* and *Henipavirus* use different host receptors than sialic acid. From our novel sequences, the PMV sequences from Heteromyid rodents don’t appear to use N-Acetylneuraminic acid based on conservation at the active site, despite being nested within the larger *Jeilongvirus*-like viruses that do mainly appear to use the N-Acetylneuraminic acid sugar motif. What is especially remarkable is that despite high levels of sequence divergence in the H/HN/G locus, phylogenetically diverse PMVs from a broad range of hosts use N-Acetylneuramic acid as a host receptor. It remains unclear whether this pattern is due to N-Acetylneuramic acid being the ancestral host receptor for all *Orthoparamyxovirinae*, or whether diverse viruses have convergently evolved to use N-Acetylneuramic acid This pattern is especially clear in the phylogenies shown in Fig. 4.

All of the novel bat and rodent PMVs reported here, and other representative sequences from *Jeilongvirus* and *Shaanvirus*, are sister to the *Respirovirus* genus at the H/HN/G locus, although for all other loci *Respirovirus* is the most divergent relative to other major genera. One possible explanation for this discordance is recombination, however recombination is thought to be extremely rare in negative-sense single stranded RNA viruses (46). Instead, the flexibility of the paramyxovirus attachment protein likely appears to be tolerant to high sequence divergence while allowing for frequent switching between different host receptors during the evolutionary history of PMVs.

## Acknowledgments

B.B.L. was supported by funding from the NSF GRFP, Society for the Study of Evolution Rosemary Award, and the University of Arizona Galileo Scholar program. Funding was provided to M.W. from the David and Lucile Packard Foundation.

